# Predictability changes what we remember in familiar temporal contexts

**DOI:** 10.1101/469965

**Authors:** Hyojeong Kim, Margaret L. Schlichting, Alison R. Preston, Jarrod A. Lewis-Peacock

## Abstract

The human brain constantly anticipates the future based on memories of the past. Encountering a familiar situation reactivates memory of previous encounters which can trigger a prediction of what comes next to facilitate responsiveness. However, a prediction error can lead to pruning of the offending memory, a process that weakens its representation in the brain and leads to forgetting. Our goal in this study was to evaluate whether memories are spared from such pruning in situations that allow for accurate predictions at the categorical level despite prediction errors at the item level. Participants viewed a sequence of objects, some of which reappeared multiple times (“cues”), followed always by novel items. Half of the cues were followed by new items from different (unpredictable) categories, while others were followed by new items from a single (predictable) category. Pattern classification of fMRI data was used to identify category-specific predictions after each cue. Pruning was observed only in unpredictable contexts, while encoding of new items was less robust in predictable contexts. These findings demonstrate that how associative memories are updated is influenced by the reliability of abstract-level predictions in familiar contexts.

## INTRODUCTION

What is past is prologue: similar to the function of autocomplete software on a smartphone, the brain learns from statistical patterns across time to generate expectations that guide future behavior. This process is essential for most of our fundamental abilities including language, perception, action, and memory. It is accomplished in part by domain-general implicit learning mechanisms – alternatively referred to as ‘statistical learning’ (Perruchet & Pacton, 2006; Turk-Browne, Jungé, & Scholl, 2005; Turk-Browne & Scholl, 2009) – that allow us to acquire long-term knowledge about the statistical structure of the world (Kóbor, Janacsek, Takács, & Nemeth, 2017; Romano, Howard Jr, & Howard, 2010). Studies of visual statistical learning have shown that observers can implicitly learn subtle statistical relationships between visual stimuli in both time (Fiser & Aslin, 2002; Fiser, Scholl, & Aslin, 2007; Schapiro, Kustner, & Turk-Browne, 2012) and space (Chun & Jiang, 1998; Fiser & Aslin, 2001; Preston & Gabrieli, 2008; Turk-Browne & Scholl, 2009). Knowledge of these statistics can build up expectations that trigger predictions about upcoming perceptual events (Turk-Browne, Scholl, Johnson, & Chun, 2010). In some situations, these predictions may be stimulus-specific (Conway & Christiansen, 2006), and in others they may be more abstract, for example, operating at a categorical level that relies on existing conceptual knowledge (Brady & Oliva, 2008).

An advantage of predicting upcoming events is to support fast and adaptive responding to the environment when those predictions are accurate. However, prediction errors (Pagnoni, Zink, Montague, & Berns, 2002; Schultz & Dickinson, 2000) also serve an adaptive role by “pruning” the memories that produced those errors, thereby reducing the likelihood of similar errors in the future. Memory pruning is an error-driven learning process in which memories that trigger mispredictions are weakened via competitive neural dynamics (Kim, Lewis-Peacock, Norman, & Turk-Browne, 2014; Kim, Norman, & Turk-Browne, 2017). Prediction error may not always lead to pruning, however. For example, the fidelity of reactivated predictions is graded, and pruning is most likely for items that are moderately, but not strongly reactivated (Kim et al., 2014). When reactivated predictions don’t get pruned, they may instead become integrated with the new event (Morton, Sherrill, & Preston, 2017; Preston & Eichenbaum, 2013; Schlichting & Frankland, 2017; Schlichting, Mumford, & Preston, 2015; Schlichting & Preston, 2015; Zeithamova, Dominick, & Preston, 2012; Zeithamova, Schlichting, & Preston, 2012). Importantly, it is unclear why certain memories are pruned and others are integrated. Therefore, a central goal of this study was to evaluate how the predictability of events influences memory updating.

To date, memory pruning has been studied only in unpredictable situations that consistently generate prediction errors. It is unknown how pruning operates in more predictable situations. Our experiences often contain hidden statistical structure that can be learned and generalized to new events. For example, existing semantic knowledge of the world can be used to link conceptually related episodic events across time to form situation-specific “schemas” to guide future behavior (Ghosh & Gilboa, 2014; Preston, Molitor, Pudhiyidath, & Schlichting, 2017; Tse et al., 2007; van Kesteren, Rijpkema, Ruiter, & Fernández, 2010; van Kesteren, Ruiter, Fernández, & Henson, 2012). If every time you walk to your neighborhood park you encounter a new dog, you may come to form a schema that parks are associated with the presence of dogs. Such a schema would allow for future predictions at an abstract level, but not for item-specific details (e.g., “I expect to see some dog at this park” but not “I expect to see that fluffy poodle named Ruby at this park”). More generally, if certain items in a sequence of events are always followed by the same *category* of item, this should allow for more abstract, categorical-based statistical learning and prediction (Brady & Oliva, 2008). Over time the brain will adapt to the predictability of these category exemplars and as a result build expectations about a category rather than any specific item (Brown & Braver, 2005; den Ouden, Friston, Daw, McIntosh, & Stephan, 2009).

In this study, we manipulated the predictability of images in different temporal contexts and evaluated the impact on subsequent recognition memory for those images. A context where it is difficult to predict which items will appear next should produce item-level prediction errors. This context should lead to pruning of the mispredicted items, and it could also be associated with strong encoding of new items because their predictive relationship with the environment is still being learned (Dayan, Kakade, & Montague, 2000). Alternatively, a context that allows for accurate categorical predictions should lead to less pruning of item-specific representations. As predictions shift to the category level, the specific items would no longer be predicted and therefore their neural representations would not be susceptible to the pruning process. However, the predictability of this context may come at the expense of detailed stimulus processing, thus leading to weaker encoding of new items (see Rommers & Federmeier, 2018).

To test these ideas, we presented participants with a continuous sequence of visual object presentations. In this incidental encoding task, observers made subcategory judgments about each object in the sequence, and memory for these items was tested later with a surprise item-recognition memory test. As a hidden rule, certain items (“cues”) appeared four times across the experiment, and all other items appeared only once. To manipulate the reliability of predictions generated by the cues, half of the cues were followed by items from *different* categories across repetitions (unpredictable context), and the other half of cues were followed by items from the *same* category (predictable context). Functional MRI (fMRI) data was collected during this encoding task, and in a separate functional localizer task, and we used multivoxel pattern analysis (MVPA; Haxby et al., 2014; Haynes & Rees, 2006; Lewis-Peacock & Norman, 2014a; Norman et al., 2006) to quantify the perception of each stimulus, and to covertly measure the automatic prediction of items following the reappearance of the cues. These trial-by-trial neural measures were then linked to item-recognition performance on the subsequent memory test. Memory pruning was evaluated at each cue repetition by assessing the relationship between neural evidence of prediction of previous items and their subsequent memory strength, whereas encoding strength was evaluated by assessing the relationship between neural evidence of perception of new items and their subsequent memory strength.

## METHODS

### Participants

Thirty healthy young adults (13 male; age, *M* = 22 yr, *SD* = 3.48, all right-handed) were recruited from the student body and campus community of the University of Texas at Austin to participate in the neuroimaging experiment. Five participants were excluded due to low classifier accuracy in the localizer task (5 *SEM* below the mean), which was an a priori criterion for applying the classifiers to the encoding data. Three more participants were excluded due to low recognition accuracy (10 *SEM* below the mean), resulting in final sample size of *n =* 22. Twenty-four additional participants (13 male; age, *M* = 23.29y, *SD* = 4.81, left-handed = 1) were recruited for a behavior-only version of the experiment. All participants had normal or corrected-to-normal vision. The study was approved by the University of Texas at Austin Institutional Review Board and informed consent was obtained from all participants.

### Stimuli

Colored pictures of common objects were used for this experiment. They were selected from six categories (with two subcategories each): famous faces (female/male), famous scenes (manmade/natural), animals (land/non-land), food (cooked/uncooked), tools (power/non-power), and vehicles (land/non-land). Object images were obtained from various resources (Morton, Zippi, & Preston, in prep.) including Bank of standardized stimuli (Brodeur, Guérard, & Bouras, 2014), and Google Images.

### Procedure

The experiment proceeded in three tasks: incidental encoding, functional localizer, and subsequent recognition memory test (Figure 2A). FMRI brain data was acquired for the encoding task (6 runs, 335s/run) and the localizer task (2 runs, 513s/run). Participants (*N* = 22, Figure 1A). Participants (*N* = 22) performed the subsequent recognition memory test either after the localizer (outside the scanner, *n* = 14) or before the localizer (in the scanner, *n* = 8). For behavior-only participants (*N* = 24), encoding and recognition phases were conducted sequentially.

**Figure 1.**
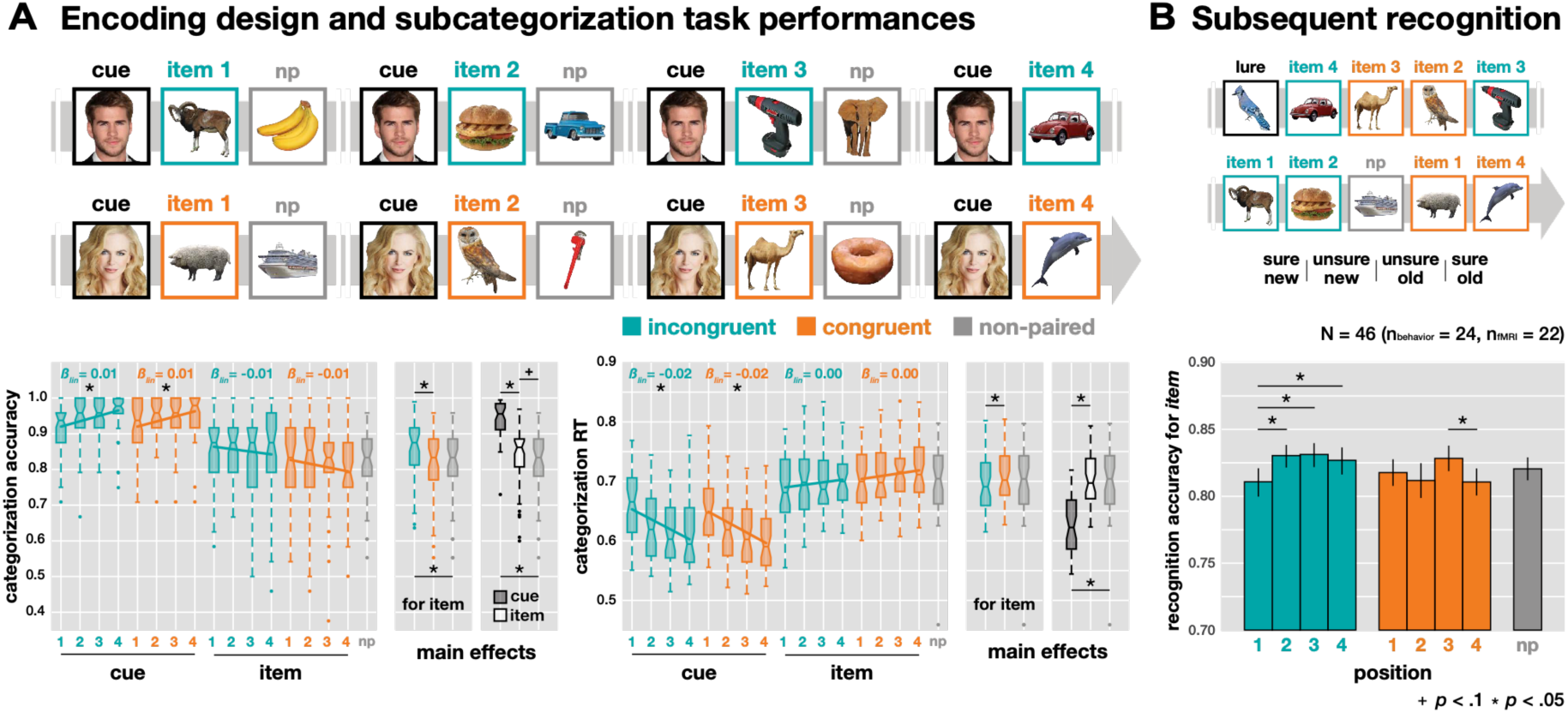
Behavioral performance. (A) Incidental encoding task with categorization accuracy and RT. Subcategory judgments for each picture in a sequence (CUE: items that were presented four total times; 1^st^/2^nd^/3^rd^/4^th^ ITEM: unique items that followed a cue in the specified order; NP: non-paired items that did not follow cues). The category for the cues was either face or scene, and there were four categories (animal, food, tool, and vehicle) for items and NP items. For main effects, Bonferroni adjusted alpha levels (α=.05/3) were applied per test. (B) Subsequent recognition task and recognition accuracy (A’) for all items, not cues, studied previously. Four options with old/new and sure/unsure were given for the response. Error bars represent SEM.

#### Incidental encoding task

Participants were shown a steady stream of images, one at a time, for the purpose of incidental encoding for a surprise subsequent memory task at the end of the experiment (Figure 1). There were three types of trials – cue, item, and non-paired item – across two conditions (congruent/incongruent) and four positions (1^st^/2^nd^/3^rd^/4^th^ for each pair). In the stream, a cue was associated with an item as a cue-item pair in hidden sequences. Each cue was associated with four different items, which made four cue-item pairs as one set. For half of the cues, all items were selected from a single category (*congruent condition*). For the other half of the cues, the items were selected from a new category each time (*incongruent condition*). There were 24 sets (96 total cue-item pairs) for each condition, and 96 *non-paired* items that were not part of a set and never directly followed a cue. The pairs from a given set were not adjacent but appeared intermingled with other sets and non-paired items (mean lag between pairs = 8 trials) within a single run. All cues appeared four times, and all other items appeared only once (Figure 1A). In each run (80 trials), there were four sets for both conditions and 16 non-paired items. Across all six runs (480 trials total), the categories (e.g., animal, food, etc.) and subcategories (e.g., land/non-land, cooked/uncooked, etc.) of items and non-paired items were counter-balanced.

As a cover task, participants were asked to make a subcategory judgment for each image using one of two buttons on a 4-button box (in the scanner) or on a keyboard (outside the scanner). The category and subcategory of the stimulus were randomized across trials, and therefore participants were required to constantly update their response mappings. To facilitate performance, we provided the two subcategory options for each stimulus (e.g., female/male for faces), which required different responses for the same category. On a trial, the stimulus displayed for 1 s on a white background box (visual angle: 21.8° × 21.8°), with empty feedback circles and text underneath the image displaying the subcategory choices, during which participants had to make a response. When the stimulus disappeared, a blank white box remained with feedback circles underneath, in which one of the circles was colored for 1 s based on performance (green: correct, red: incorrect, yellow: missed). The inter-trial interval (ITI) was pseudo-randomly jittered at 2, 3, or 4 s, and there was no systematic bias of ITI distributions across conditions and positions for both cues and items (two-way ANOVA, *Fs*(3, 63) < 1.33, *Ps* > .272). Moreover, there was no significant effect of jittered ITI on subsequent memory (one-way ANOVA, *F*(2, 42) = 0.73, *P* = 0.490).

Either faces or scenes, but not both, were used as cue stimuli for each participant (*N* = 13/22 fMRI, and *N* = 12/24 behavioral participants had face cues). The non-selected category was not used for the encoding task for that participant. We isolated these two categories as cues based on their superior classification accuracy in ventral temporal cortex (face, scene; *M* = 0.82, *SE* = 0.09), and selected the other four categories with similar decodability as items (*M* = 0.57, *SEM* = 0.10), based on a separate pilot sample (*N =* 3) on the localizer task (see Figure 2B). It was necessary to use unbiased visual categories for items in terms of decodability because the decoding strength of items across conditions was our main measurement. We chose famous people and famous places to facilitate recognition of the cues, which in turn should facilitate the generation of context-based predictions when the cues repeated. The other four categories (animals, food, tools, vehicles) were used for the stimuli that appeared (only once) as items following a cue or as non-paired items. Participants practiced the task before scanning with a separate set of images until they reached a criterion of 80% accuracy for the subcategory judgment task. Categorization performance was calculated with accuracy and RT of the responses, and a simple linear regression was applied to track the performance changes across repetitions for trial type and condition.

#### Subsequent recognition memory test

In this phase, the participants were given a surprise memory test for the objects that they saw in the encoding task. All objects used for items and non-paired items (288 old; 96 items for each incongruent, congruent, and non-paired condition) and 96 novel lures were tested in a random order (Figure 1B). Participants made a recognition judgment using a 4-point scale: 1 = sure new, 2 = unsure new, 3 = unsure old, and 4 = sure old. Only “sure old” responses were treated as hits (Kim et al., 2014; Lewis-Peacock & Norman, 2014a), and we calculated memory sensitivity using A-prime (*A*’; Stanislaw & Todorov, 1999). A subset of participants (*N* = 8/22) took the memory test right after the encoding task in the scanner prior to the localizer phase to minimize any possible memory interference from stimuli in the localizer. However, there was no observed impact of task order on memory performance (*F*(1, 20) = 1.137, *P* = .299). We assessed statistical reliability of the subsequent memory results for the fMRI (n = 22) and behavioral group (n = 24) separately, and also for the combined data from both groups (N = 46 total).

#### Functional localizer

Participants performed a one-back task with six categories of images: face, scene, animal, food, tool, vehicle (Figure 2A). These stimuli were unique to the localizer and were never shown again. Each image was presented for 1.5 s on a white background box followed by an inter-trial interval for 0.5 s in which only the white background box remained on the screen. Stimulus display parameters were similar to the encoding task. However, rather than making a subcategory judgment, participants responded “same” if the object matched the previous object or “different” otherwise (on average, there was 1 repeat every 5 trials). Responses were to be made within 1 s, and visual feedback was given using the color of the frame of the background box (green: correct, red: incorrect) immediately after the response, or after stimulus offset if no response was made. Stimuli were blocked by category with 10 trials per mini-block, lasting 20 s, and 6 mini-blocks (6 categories × alternate subcategory across blocks) per block, followed by 6 s of blank inter-block interval. There were two fMRI runs of the localizer task, each with 4 blocks (24 mini-blocks) presented in randomized order. Fifteen seconds were added to the end of each run to account for hemodynamic delay on the last trial. To verify the accuracy of the classifier, the one-sample *t* test was conducted for each category.

### Data acquisition

The Psychophysics Toolbox (http://psychtoolbox.org) was used to run experiments. Neuroimaging data were acquired on a 3.0-T Siemems Skyra MRI (Siemens, Erlangen, Germany) with a 64-channel head coil. High-resolution anatomical images were collected for registration from a T1-weighted 3-D MPRAGE volume (256 × 256 × 192 matrix, 1 mm^3^ voxels). A gradient-echo echo planar imaging sequence was applied for functional images with following parameters: TR = 1 s, multiband factor = 4, TE = 30 ms, 63° flip, 96 × 96 × 64 matrix, 2.4 mm^3^ voxels, 56 slices, no gap.

### Preprocessing

FSL (http://fsl.fmrib.ox.ac.uk) was used to preprocess the fMRI data. Functional volumes were corrected for motion, aligned to the mean volume of the middle run, temporal high-pass filtered (128s) and detrended. Timepoints with excessive motion were removed (framewise displacement, threshold = 0.9 mm (Power et al., 2014); *M* = 6.7 TRs removed, *SD* = 14.5).

### Region-of-interest

We were interested in bilateral ventral temporal cortex (VTC; Grill-Spector & Weiner, 2014; Haxby et al., 2001) in which high-level visual categories are predominantly represented (Figure 2A). This region of interest (ROI) was anatomically delineated in standard space by combining bilateral temporal occipital fusiform cortex and posterior parahippocampal gyrus from the Harvard–Oxford cortical atlas. This was converted to native space and resampled to functional resolution for each subject using the transformation matrix derived from registration of that subject’s data to standard space. This size of this ROI across subjects was *M* = 4,266 voxels, *SD* = 283.

We focused on VTC rather than default mode network (DMN) or hippocampus (Kim et al., 2017; Long, Lee, & Kuhl, 2016; Schapiro et al., 2012), which were examined when association learning was stronger with explicit association learning or multiple repetitions of implicit learning procedures. We applied the same ROI from the previous study that also used one-time incidental learning (Kim et al., 2014). In separate analyses with DMN and hippocampus ROIs, we found that only face and scene categories were successfully decoded from the localizer data (DMN: *M* = .52, *SEM* = .03, hippocampus: *M* = 0.47, *SEM* = 0.33), which were used as predicted items in these prior studies. However, there was poor classification accuracy (less than half as good) for the other four item categories in these ROIs (DMN: M = .23, SEM = .01, hippocampus: M = 0.22, SEM = 0.01), and therefore these regions were not interrogated further.

### Classification Analyses

The Princeton Multi-Voxel Pattern Analysis Toolbox (www.pni.princeton.edu/mvpa) was used for multivoxel pattern classification using L2-regularized, non-multinomial (one-vs-others, for each category) logistic regression. Classifiers were trained separately for each participant using localizer data in bilateral ventral temporal cortex. Regressors for all seven categories (face, scene, animal, food, tool, vehicle, rest) were shifted by 4 s to adjust for hemodynamic lag. To validate classifier performance, cross-validation was performed across the two runs of localizer data. This was done 22 times with different penalties (from 0 to 1000) to find the optimal penalty for each participant (*M* = 156, *SD* = 244). Prior to classification, feature selection was performed for each training set using a voxel-wise ANOVA across all categories and timepoints (threshold: *P* = 0.0001) with the regressor modeled with a mini-block boxcar and shifted forward 4 s to account for hemodynamic lag. The selected voxels were used to train and test the classifier (19.2% of the original voxels; *M* = 820 voxels, *SD* = 237). Across subjects, classifier performance was reliably above chance for each category (*M* = 0.58, *SEM* = 0.02, chance level = 0.14, Figure 2B). Data from both localizer runs were then used to re-train the classifiers which were then applied to data from the encoding task (Figure 2A). This produced classifier evidence scores (from 0 to 1) for each category at every timepoint in the encoding task. These scores reflect the likelihood that a test sample of brain activity contains a representation of a given category. The same individualized penalty derived from the cross-validation of the localizer data was used, and a new feature-selected mask was computed (26.6% of the original voxels; *M* = 1136, *SD* = 264). The *perception strength* of each object during the encoding task was defined as the average classifier evidence for the object’s category from its onset until the onset of the next stimulus (i.e., 1 s of display plus 2, 3, or 4 s of ITI, depending on jitter, shifted forward 4 s to account for hemodynamic lag; *prediction window*, Figure 3A). On cue repetitions, the *prediction strength* for the item that previously followed the cue was defined as the average classifier evidence for that item’s category during the perception time window for the cue (*perception window*, Figure 3A). Note that to minimize influence from the onset of the next item, but to keep the window size equal to the perception window, the prediction window began 1 s prior to cue onset, which was adopted from the previous study (Kim et al., 2014), and then shifted 4 s forward to account for hemodynamic lag. In a separate analysis, we applied fixed 3 s windows for perception and prediction from the onset of each trial and found no significant deviations in the main findings (see Results).

Before the main analyses, which linked the classifier evidence strength to memory, we compared both prediction and perception evidence distributions from category classifier across conditions and positions. Classifier evidence was averaged for each process × condition × position within subject. Across all positions (i.e., 2^nd^ – 4^th^ for prediction, 1^st^ - 4^th^ for perception), repeated-measure two-way ANOVAs were conducted for condition × position within each process, and then three-way ANOVAs were applied for process × condition × position for the final three positions (Figure 3A). Finally, the classifier evidence scores for the target category (e.g., the perceived or predicted category) were compared to the mean of the three non-target categories with a three-way ANOVA for condition × position × target (target/non-target).

### Trial-level modeling

In an attempt to improve decoding sensitivity for predictions in this fast event-related design, we modeled category-level beta estimates for the perception of each stimulus in the encoding task (General Linear Model, GLM, utilizing a hemodynamic response function) to remove the evoked activity from cue presentations. Then we modeled trial-specific beta estimates for the predictions from the residuals of this analysis (GLM including a regressor for that trial + another regressor for other trials; Mumford, Turner, Ashby, & Poldrack, 2012). We applied the same classifiers trained on the localizer data with shifted regressors to decode the “prediction betas” from the encoding task. The results obtained from this analysis were qualitatively similar to the results obtained using the unmodeled, shifted regressors, which we chose to report here.

### Linking neural data to behavior

Binary logistic regression analysis was used to examine the impact on subsequent memory from prediction strength and perception strength during the encoding task (Figure 3B). The result of each of these analyses is a coefficient estimate (β) of the relationship between the given neural evidence and the memory outcomes. To increase our ability to detect trial-specific effects with our small sample size, we pooled data from all subjects and then performed bootstrap resampling to evaluate the population reliability of the result (Efron, 1979; Fisher & Hall, 1991). On each bootstrap iteration (of 1,000 total), we sampled randomly (with replacement) a collection of participants’ data to match the size of our experimental sample (*N =* 22). There were eight regressions conducted for perception strength and subsequent memory (incongruent/congruent × 1^st^/2^nd^/3^rd^/4^th^ positions), and six regressions conducted for prediction strength and subsequent memory (incongruent/congruent × 2^nd^/3^rd^/4^th^ positions). Statistical significance was calculated with a non-parametric test across bootstrap iterations, evaluating the stability of an effect of interest by calculating the proportion of iterations in which the effect was found. Lastly, to verify that the effects were not arising from variance across participants but from within-subject variance, we repeated the main analyses using standardized (z-scored) classifier evidence for each participant to remove subject-specific effects (Kim et al., 2014). Results from the main analyses were qualitatively similar and confirmed. Across repetitions, linear regression analyses were conducted on the binary logistic regression results (β) across positions for each condition and process, and bootstrapped for the statistical significance test. Then, a repeated-measure two-way ANOVA for condition × process was conducted on the coefficient of linear regressions (β_lin_) across positions, and the statistically reliability was assessed via bootstrapping.

### Pruning effect searchlight

We examined the whole brain to exhibit pruning effect with searchlight analysis. First, we limited the voxels (i.e., center of sphere, radius = 2mm) in which the classification accuracy for item-categories with localizer data was higher than chance level, 0.14. Then, we narrowed down the voxels that had a negative relationship (β) between prediction evidence and memory strength (the signature of memory pruning) and had more negative relationship for the incongruent than the congruent condition. The β-values from the selected voxels were then resampled (6 × 6 × 6 mm^3^), normalized into MNI space. For group-level analysis, only the voxels that selected for more than twenty participants and whose β-values had a significant effect across participants were collected (voxel-wise one-sample *t* test, α = .01, FDR correction for multiple comparisons). The final voxels were cluster corrected (extent voxels = 10) and smoothed (12mm FWHM) as a binary mask (Figure 6).

## RESULTS

Note that for all behavioral results, we report combined results (*N* = 46). For the subsequent memory, we also report results separately for fMRI participants (*n* = 22) and behavior-only participants (*n* = 24) who performed the same task outside the scanner (see Methods).

### Encoding task performance

Participants were shown a continuous stream of images, one at a time, for the purpose of incidental encoding for a surprise subsequent memory task at the end of the experiment (Figure 1). As a cover task, participants were asked to make a subcategory judgment for each image. The subcategory of the stimulus was pseudo-randomly assigned across trials, which made the judgment task unpredictable. Thus, participants were required to maintain their attention to each stimulus and update their response mapping to make an accurate judgment within the 1s response-window. Subcategory judgments were fast (*M* = 0.67 s, *SEM* = 0.01) and accurate (*M* = 0.87, *SEM* = 0.01, Figure 1A). Performance differed across conditions (incongruent, congruent), positions (1^st^, 2^nd^, 3^rd^, 4^th^), and non-paired items for both trial types (cue, item) in both accuracy and RT (one-way ANOVA, omnibus *Fs* > 35.90, *Ps* < .001). In a pair, a cue and an item were sequentially associated, and non-paired items were randomly distributed between cue-item pairs without associations. Responses for cues were faster (0.62 s) and more accurate (0.94) than for items (0.70 s, 0.83) and non-paired items (0.70 s, 0.82; pairwise *t* test, all *Ps* < .001, significant after Bonferroni correction for 3 comparisons), suggesting that repeated encoding enhanced subcategorization performance. Across repetitions, a simple linear regression (coefficient estimate: β_lin_) shows that the accuracy to cues increased (all β_lin_ > 0.01, *Ps* < .01) and the RTs decreased (all β_lin_ < −0.01, *Ps* < .001), with no difference between conditions (condition × position two-way ANOVA for cues, *Fs* < 0.97, *Ps* > .41). Categorization performance on the items did not change significantly across repetitions in both conditions (*Ps* > .05).

**Figure 2.**
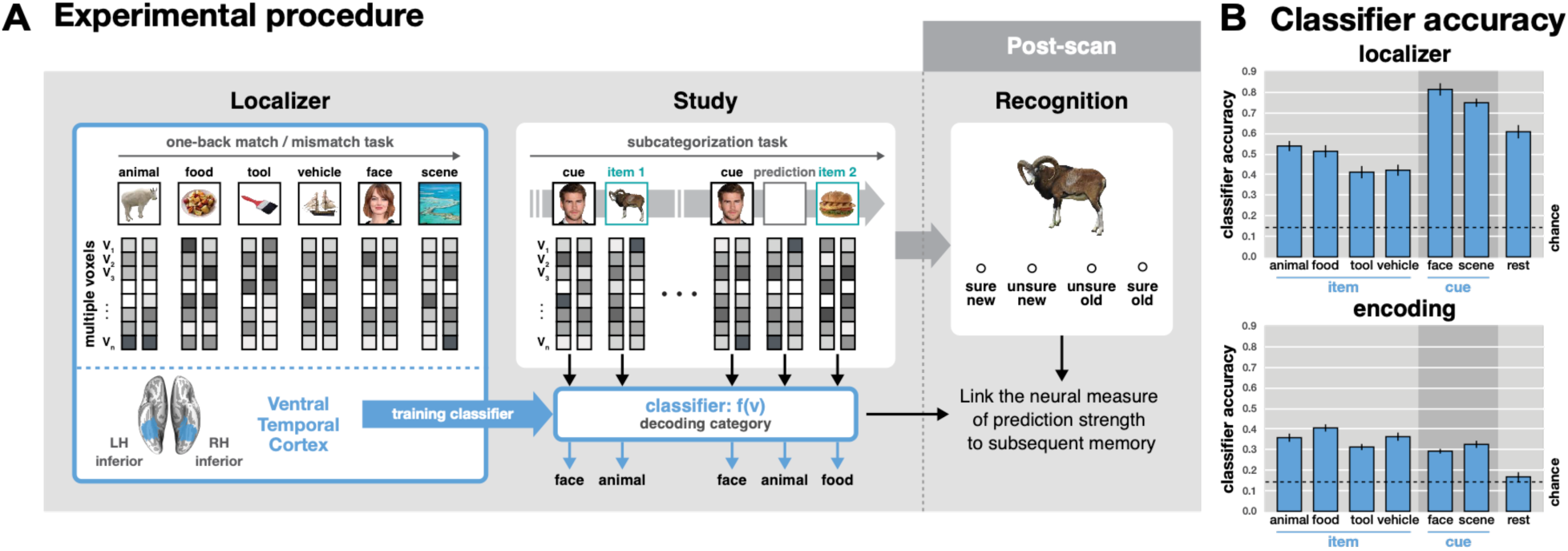
Experimental procedures and fMRI pattern classifier accuracy. (A) Experimental procedures. Pattern classifiers were trained on fMRI data from the localizer task to identify category-specific patterns of brain activity in ventral temporal cortex mask. These classifiers were then applied to the encoding task, and the neural results were linked to subsequent memory scores in the surprise recognition test. (B) Classifier accuracy. Pattern classification was successful for all categories in the localizer task (chance = 0.14). These classifiers, trained on all localizer data, successfully identified the category of each item presented in the encoding task.

There was a significant interaction of condition × trial type (cue/item) on both RT and accuracy (both *Ps* < .001), with no difference for cues, but with both faster and more accurate responses for items in the incongruent condition (0.71 s vs. 0.70 s, 0.81 vs. 0.85; both *Ps* < .001). There was no three-way interaction of condition × position × trial type (cue/item) on either RT or accuracy. Overall the behavioral metrics on the encoding task indicate that participants were properly engaged in the task, and performance differences between the incongruent/congruent conditions demonstrate that they were sensitive to this manipulation.

### Subsequent recognition memory

Memory for all items was tested in a surprise recognition test at the end of the experiment (Figure 1B; Table 1). In the incongruent condition, there was worse memory for the 1^st^ items (*M* = 0.81, *SEM* = 0.01) compared to the other items (2^nd^/3^rd^/4^th^, *M*s > 0.83, *SEMs* < 0.01; pairwise *t* tests, all *Ps* < .040). In the congruent condition, there was worse memory for the 4^th^ items (*M* = 0.81, *SEM* = 0.01) compared to the 3^rd^ items (*M* = 0.83, *SEM* = 0.01; *t*_45_ = 2.35, *P* = .023). There were no significant interactions between conditions or compared to baseline (non-paired) items. Note that our main analyses focused on how neural signals related to memory consequences on a trial-by-trial basis, rather than the overall memory performance itself.

**Table 1.** Subsequent recognition performance. (A) Recognition accuracy. Mean for each position and condition with 95% CI in the next parenthesis. (B) Pairwise *t* test. All significant comparisons (*P* < .05) within each condition were included across data sets and white cells indicate non-significant comparisons.

| Study | incongruent |  |  |  | congruent |  |  |  | NP | Total |
| --- | --- | --- | --- | --- | --- | --- | --- | --- | --- | --- |
|  | 1st | 2nd | 3rd | 4th | 1st | 2nd | 3rd | 4th |  |  |
| Combined (N = 46) | .811 (.021) | .831 (.016) | .832 (.018) | .827 (.020) | .818 (.019) | .812 (.026) | .829 (.019) | .811 (.021) | .821 (.017) | .848 (.017) |
| Behavior (N = 24) | .837 (.021) | .851 (.016) | .865 (.018) | .862 (.020) | .838 (.019) | .845 (.026) | .855 (.019) | .833 (.021) | .846 (.017) | .821 (.017) |
| fMRI (N = 24) | .783 (.026) | .809 (.020) | .795 (.025) | .789 (.027) | .796 (.028) | .776 (.041) | .080 (.027) | .787 (.032) | .793 (.023) | .792 (.023) |

**Table 1.** Subsequent recognition performance.
| Study | incongruent |  |  | congruent |
| --- | --- | --- | --- | --- |
|  | 1st vs. 2nd | 1st vs. 3rd | 1st vs. 4th | 3rd vs. 4th |
| Combined (N = 46) | $t_{45} = -2.84, P = .007$ | $t_{45} = -2.89, P = .006$ | $t_{45} = -2.12, P = .040$ | $t_{45} = 2.35, P = .023$ |
| Behavior (N = 24) | $t_{45} = -1.44, P = .164$ | $t_{45} = -2.56, P = .018$ | $t_{45} = -2.36, P = .027$ | $t_{45} = 2.83, P = .010$ |
| fMRI (N = 24) | $t_{45} = -2.60, P = .017$ | $t_{45} = -1.41, P = .172$ | $t_{45} = -0.60, P = .554$ | $t_{45} = 0.96, P = .346$ |

### fMRI pattern classification

The encoding task data were analyzed by fMRI pattern classifiers trained, separately for each individual, on category-specific data from an independent localizer task (Figure 2A, see Methods). The stimuli used in the localizer task were representatives of the same categories but were separate exemplars from those used in the encoding task. The localizer consisted of a one-back working memory task with six categories of images (face, scene, animal, food, tool, and vehicle). Within the localizer data, we verified that brain activity patterns associated with processing each stimulus category were reliably differentiated in ventral temporal cortex (*M* = 0.58, *SEM* = 0.02, chance = 0.14 for 6 stimulus categories + rest; one-sample *t* test, *Ps* < .001, Figure 2B), using independent training and testing sets with cross-validation analysis. Decoding accuracy for the cue categories (face and scene) was reliably higher than for the other four categories (animal, food, tool, and vehicle; paired *t* test, *t*_21_ = −15.33, *P* < .001), which is unsurprising given the ubiquitous use of faces and scenes in the literature (see Grill-Spector & Weiner, 2014). Pattern classifiers were then re-trained on all data (2 runs) from the localizer task and applied to the encoding task to decode every timepoint in the experiment.

As a sanity check, we verified that the category of all items presented in the encoding task was being accurately classified (*M* = 0.32, *SEM* = 0.01, chance = 0.14, *Ps* < .001, Figure 2B). However, for cues (which appeared four times each), the classification accuracy was *lower* than for items (which appeared only once; paired *t* test, *t*_21_ = 3.24, *P* = .004). This was true in both the incongruent condition (*t*_21_ = 2.75*, P* = .012) and congruent condition (*t*_21_= 2.95, *P* = .008). This relationship is a reversal from the results in the localizer data described above where the cue categories (faces and scenes) were classified with greater accuracy than the other four categories. This reduction in decoding accuracy for the cues in the encoding task likely reflects the co-mingling of cue processing and automatic predictions of items from other categories triggered by the reappearance of the cue. This possibility will be addressed further in the Discussion.

To quantify both the perception of viewed items and the prediction of subsequent items in the encoding task, we defined two neural metrics using the fMRI pattern classifiers. The “*perception strength*” for each item was defined as the amount of classifier evidence for that item during its presentation (Figure 3A). There were no differences in mean perception strength for items across the four sequence positions (1^st^, 2^nd^, 3^rd^, 4^th^) and two conditions (congruent, incongruent; repeated measure two-way ANOVA, condition (main effect): *F*(1, 63) = 0.04, *P* = .835, position: *F*(3, 63) = 1.73, *P* = .171, interaction: *F*(3, 63) = 0.32, *P* = .811). The “*prediction strength*” for an item was defined as the amount of classifier evidence for that item when its cue reappeared later in the sequence. For example, in the example sequence in Figure 3A (“*Liam Hemsworth*, *ram, …, Liam Hemsworth*, *sandwich*”), the prediction strength for “*ram*” would be the amount of animal-category classifier evidence detected when “*Liam Hemsworth*” appeared the next time (prior to the appearance of “*sandwich*”). There were no differences in mean prediction strength across the three repetitions of the cue (2^nd^, 3^rd^, 4^th^ positions) and two conditions (congruent, incongruent; condition: *F*(1, 42) = 0.01, *P* = .942, position: *F*(2, 42) = 0.05, *P* = 0.955, condition × position: *F*(2, 42) = 1.05, *P* = .360). The distributions of scores, which were concatenated across 2^nd^ to 4^th^ positions, for both perception strength and prediction strength are shown separately for each condition in Figure 3A. Perception strength for new items was reliably higher than the prediction strength for the expected items in both conditions and all positions (*M* = 0.70 vs. *M =* 0.52; three-way ANOVA, process: *F*(1, 42) = 244.41, *P* < .001, with no other significant effects). To assess the specificity of these neural measures, the classifier evidence values for the category of target items were compared with those of non-target categories. The perception evidence was significantly higher for target categories than non-targets categories across conditions and positions (three-way ANOVA, target: *F*(1, 63) = 272.88, *P* < .001), while no such differences were found for prediction evidence (all *Ps* > .164). Note that other signals from presented stimuli might have been also represented while prediction was generated, which could weaken the prediction selectivity. In a separate analysis, we decoded predictions from the prediction betas (see Methods) in which the prediction signals were isolated from the perception signals associated with the presentation of each stimulus. The decoding accuracy for the predictions was significantly higher than chance level, 0.25 (*t*_21_ = 3.67, *p* = 0.0014). The classifier evidence for the target predictions were also significantly higher than non-targets (*t*_21_ = 4.82, *p* < 0.001) when excluding the following items from the non-targets (i.e., in the incongruent only) to minimize any influence of following items. Also note that there were no differences in correlation between presented cue signals, which we modeled out, and prediction signals across conditions and positions (2-way ANOVA, *F*(2, 42) = 0.17, *P* = .845), verifying that removing the perception-related signals did not introduce any systematic bias into the prediction measurements. Importantly, our main interests were if prediction evidence for the target was selectively linked to subsequent memory, compared to that for non-targets, which reflects a prediction-based pruning effect. We report this selectivity in the section for control analyses later.

Our next step was to link these neural measurements of prediction and perception with subsequent memory outcomes. Note that for all items that appeared in the 1^st^, 2^nd^, or 3^rd^ positions of a cue sequence, we will be assessing *two* contributions to each item’s subsequent memory outcome: its perception strength in position N, and its prediction strength in position N+1. Memory for items in the 4^th^ position had a contribution only from perception strength, because the cue did not repeat a 5^th^ time. We will first address these two contributions separately, and then together.

**Figure 3.**
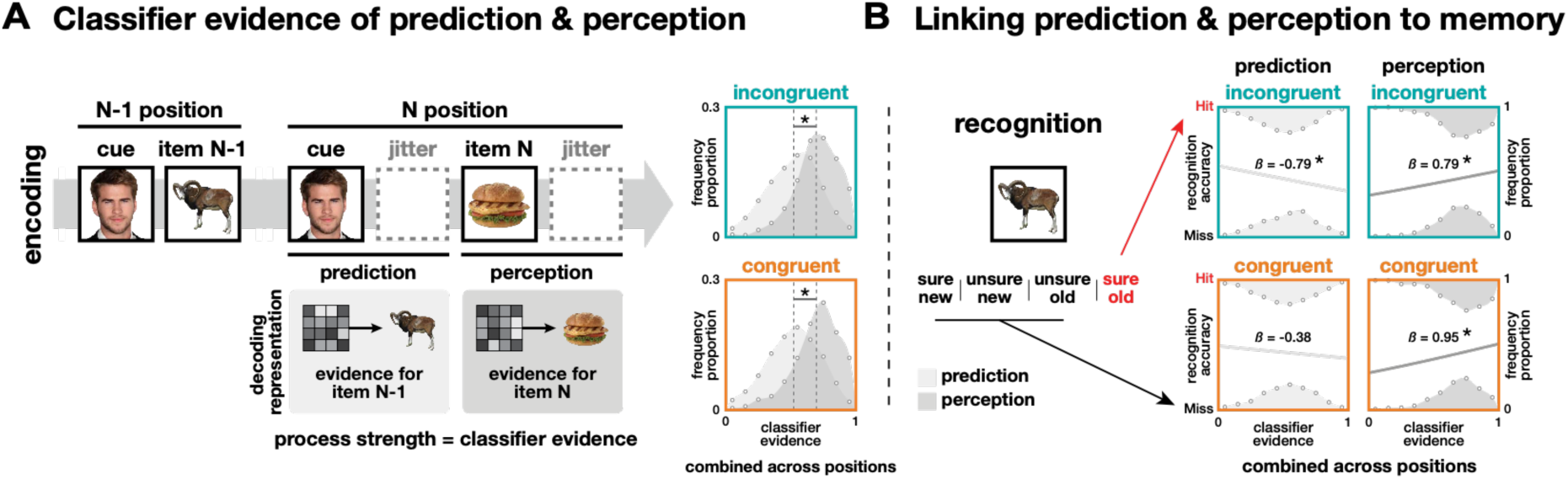
Predicting subsequent memory from brain activity during encoding. (A) Classifier evidence of prediction (item N-1) and perception (item N) in the N position (B) linking to memory performance of those items. (A) The distributions of classifier evidence for prediction (light gray) and perception (dark gray) combined across positions are shown in both conditions. (B) Logistic regression results (coefficient estimate: β) linking classifier evidence and recognition accuracy are shown separately predicted items and for perceived items combined across positions. Statistics are based on bootstrap analyses with 1,000 iterations.

### Prediction and subsequent memory

To evaluate our main hypothesis that the reliability of context-based predictions impacts pruning of associative memories, we used logistic regression to link our neural measures of prediction with subsequent recognition memory from the end of the experiment. Evidence of pruning exists when stronger predictions are associated with worse memory for the mispredicted items (see Kim et al., 2014). Figure 3B shows evidence consistent with pruning in the incongruent condition, such that stronger predictions were indeed associated with worse memory for the mispredicted items (β = −0.79, *P <* .001). This relationship was found in each repetition of the cue (2^nd^/3^rd^/4^th^ positions, *Ps* < .045), with no change across the positions (β_lin_ = 0.12, *P* = .323, Figure 4A). Consistent with our hypothesis, however, no evidence of pruning was found in the congruent condition (β = −0.38, *P* = .119, Figure 3B), where prediction strength was unrelated to subsequent memory for the mispredicted items. This relationship was absent for each repetition of the congruent cues (Figure 4B). Across all positions, there was no significant difference in the prediction-memory relationship between conditions (β_inc_ = −0.79 vs. β_con_ = −0.38, *P* = .161, Figure 4B). However, there was a statistical trend that it was stronger in the last position for the incongruent condition (β_inc_ = −0.74 vs. β_con_ = 0.23, *P* = .082, Figure 4). Although prediction strength in the congruent condition was unrelated to subsequent memory for the mispredicted item, it was, however, related to subsequent memory for the *new item*. Stronger predictions from a congruent cue were associated with worse subsequent memory for the next item that appeared (β = −0.64, *P* = .008). This was not true for incongruent cues (β = 0.01, *P* = .470; *P* = .063 vs. congruent cues). This result suggests that congruent cues, which afforded categorical predictions, biased attention toward category-general features, and away from item-specific features during encoding of the next items in the sequence. This idea will be developed further in later sections.

**Figure 4.**
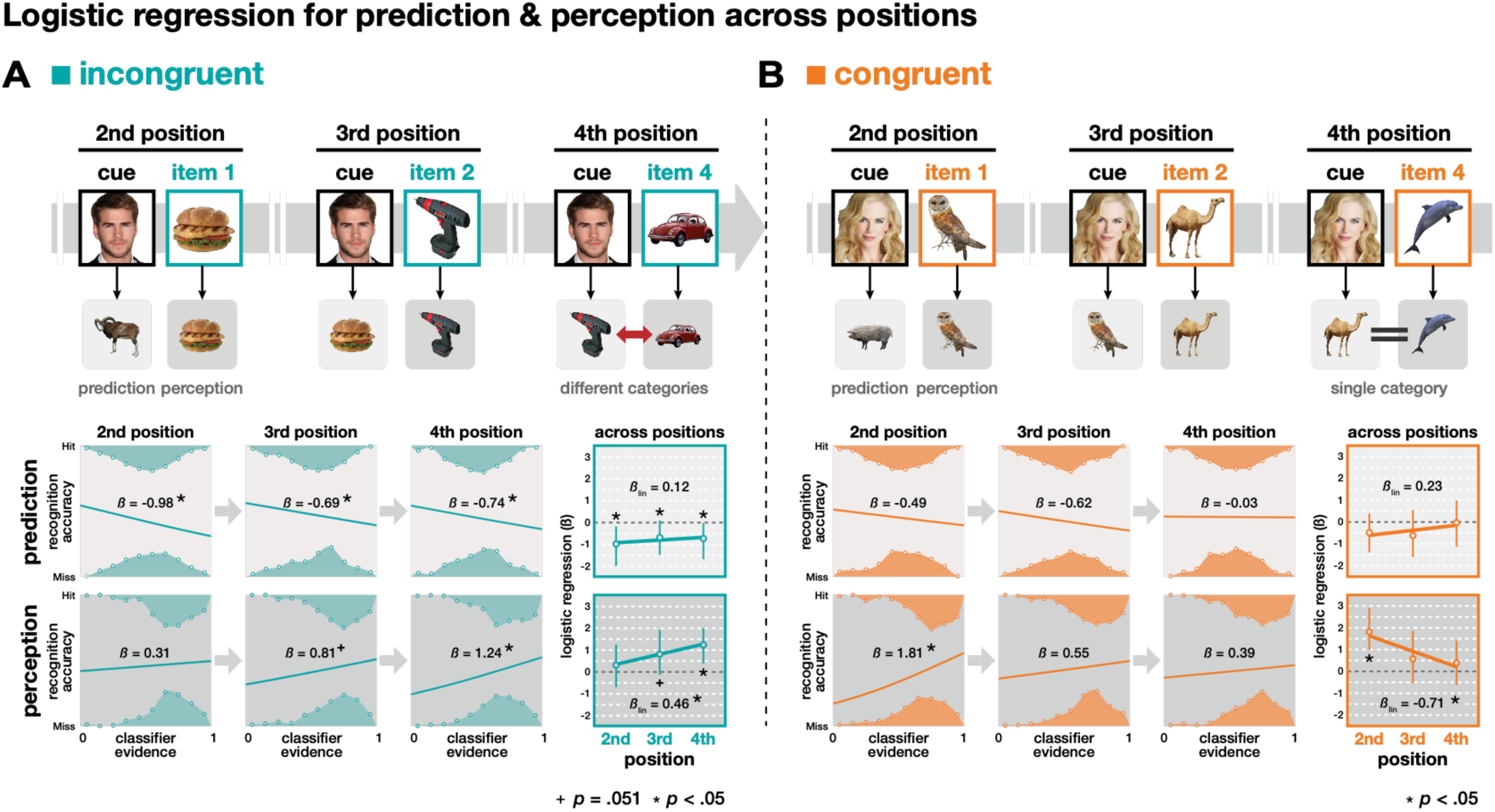
Changes across time in the links between perception, prediction, and subsequent memory in the (A) incongruent and (B) congruent condition. Hypothesized mental representations for prediction and perception across given trials (top). Logistic regression results (coefficient estimate: β) linking classifier evidence and recognition accuracy are shown separately for prediction (middle) and perception (bottom) for each position and across positions in both the incongruent and congruent conditions. Statistics are based on bootstrap analyses with 1,000 iterations. Error bars represent 95% CI.

To examine when prediction information emerged during a cue presentation, we linked subsequent memory with prediction strength measured separately from three time periods of each cue: *baseline* (3 s before the cue), *cue* (3-5 s during the cue), and *item* (3-5 s after the cue when the new item appeared, Figure 5A). As expected, there was no relationship between prediction strength and subsequent memory in either condition during the baseline period, a time at which there would be no basis for a prediction. As expected, no relationship emerged during the cue period or item period in the congruent condition (*P*s > .270). However, in the incongruent condition, the negative relationship between prediction strength and subsequent memory emerged during the cue period and persisted through the item period (*P*s < .024, Figure 5A).

**Figure 5.**
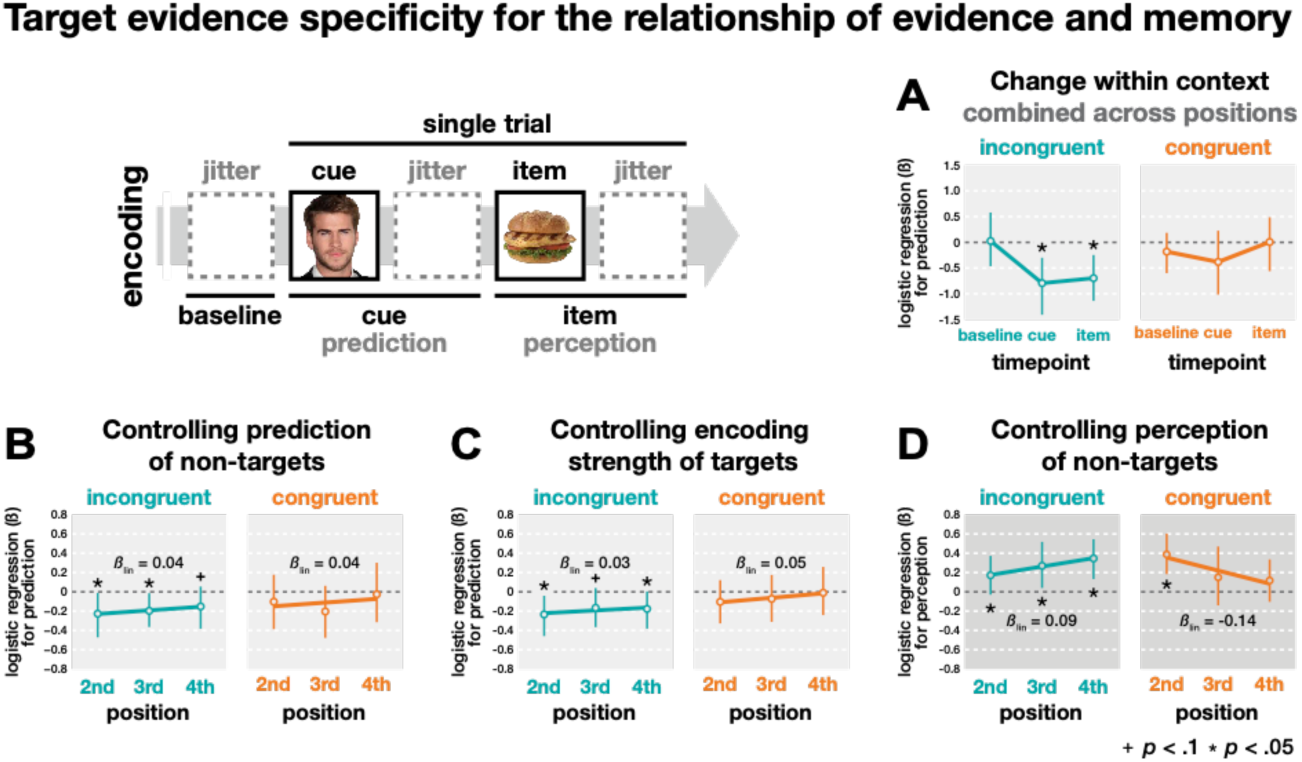
Control analyses for the specificity of the target category in the relationship between prediction/perception strength and subsequent memory across positions. (A) The relationship between prediction strength and subsequent memory is shown for different time windows *within* a cue presentation (baseline, cue, item), and averaged across the final three repetitions of each cue. (B) Controlling for the prediction evidence for non-target categories. (C) Controlling for the perception strength of the predicted item when it was encoded on its previous position following cue. (D) Controlling for the perception evidence for non-target categories. Statistics are based on bootstrap analysis with 1,000 iterations. Error bars represent 95% CI.

**Figure 6.**
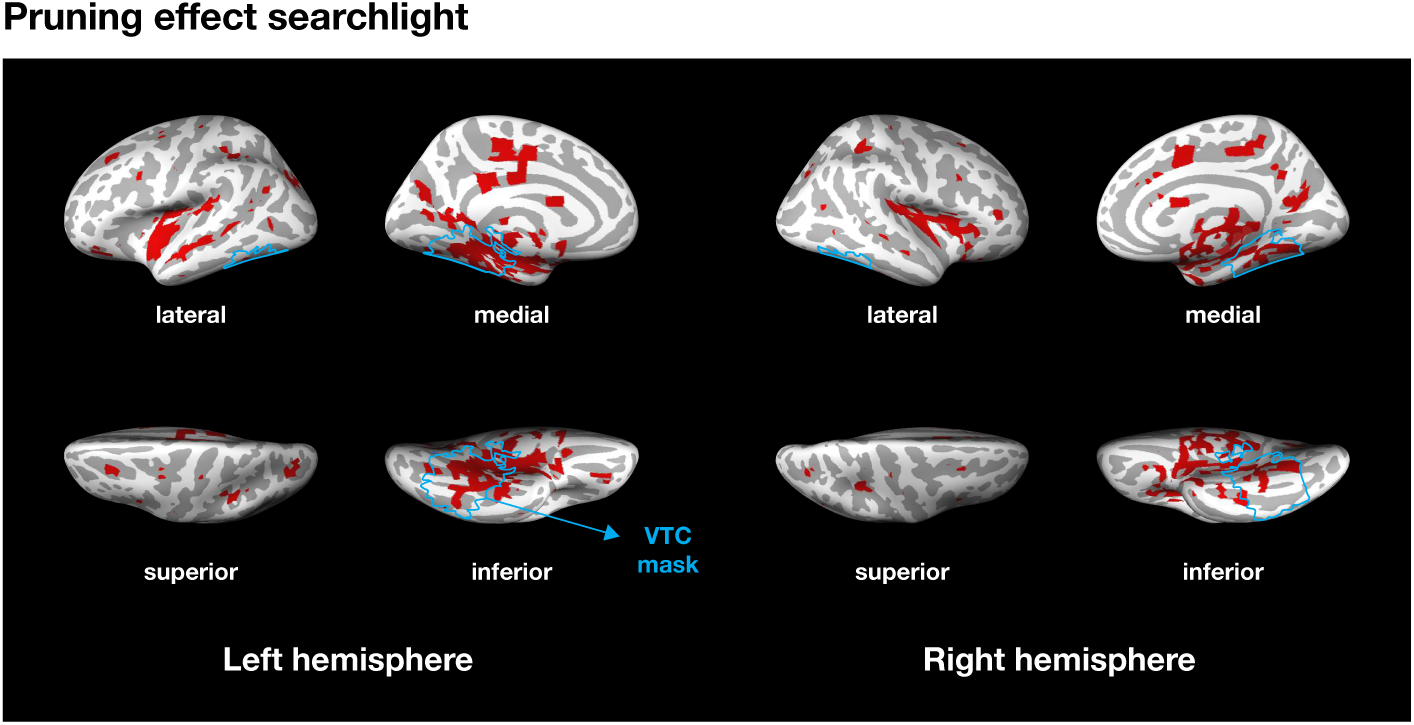
Pruning effect searchlight. Selecting voxels that showed pruning effect: a negative relationship (β) between prediction evidence and memory strength and more negative relationship for the incongruent than the congruent condition. The blue line indicates VTC ROI in 6 × 6 × 6 mm^3^ space.

### Perception and subsequent memory

To evaluate our hypothesis that increased predictability of items in a familiar context will decrease perceptual encoding of novel stimuli (or specific items), we again used logistic regression to link our neural measures of perception strength with subsequent recognition memory. In the incongruent condition, when new items were unpredictable, we found that on average stronger perception strength for an item was associated with better memory for that item (β = 0.79, *P* = .005, Figure 3B). This relationship strengthened across repetitions (β_lin_ = 0.46, *P* = .002) and was strongest by the final (4^th^) repetition (β = 1.24, *P* = .004, Figure 4A). This indicates that new items in this condition were being actively encoded because their neural index of perception strength was directed related to their subsequent memory. The influence of perception strength on memory for earlier items could have been diminished by the contribution from the (mis)prediction of those items during the next cue repetition (we will address this more in the next section). This could not be the case for the final items.

For the congruent condition, we found a similar result that, on average, stronger perception was associated with better subsequent memory (β = 0.95, *P* = .001, Figure 3B). However, this relationship weakened significantly across repetitions (β_lin_ = −0.71, *P* = .006) and disappeared altogether by the final (4^th^) position (β = 0.39, *P* = .215, Figure 4B). There was a significant interaction of condition (incongruent, congruent) on the perception-memory relationship. For earlier items, the influence of perception on memory was stronger for congruent items compared to incongruent items (1^st^ and 2^nd^, *Ps* < .027), but this relationship was reversed for the final items (4^th^, *P =* .046). Across the three repetitions, the shift in perception-memory relationship differed significantly between the two conditions (linear: β_inc_ = 0.46 vs. β_con_ = −0.71, *P* = .001). Our primary analyses focus on items in the final three repetitions, but these interaction results for perception and memory hold when including 1^st^ position items as well (linear: β_inc_ = 0.30 vs. β_con_ = −0.51, *P* = .002). Finally, there was a significant interaction of condition × process (prediction/perception) on the shift in brain-behavior relationships across repetitions (*F*(1, 1999) = 9053.83, *P* < .001) indicating that the anticipation and processing of new items impacted memory differently across time in predictable vs. unpredictable contexts.

Together, these results indicate that new items in the congruent condition, whose category could be predicted from prior statistics, were encoded less well after multiple repetitions of their context than the items in the incongruent condition. This is consistent with the idea that statistical learning at the categorical level was gradually stabilized across multiple repetitions of the context, which led to new items being also encoded at the categorical level at the expense of item details. This does not imply that these items were being ignored, however, as they could be processed sufficiently to provide successful recognition performance even without a link between perception strength and memory. We will now discuss various control analyses that rule out possible confounds and alternative explanations of our results.

### Controlling for potential confounding factors

Control analyses confirmed that the negative relationships of prediction strength and memory in the incongruent condition (i.e., memory pruning) were specific to classifier evidence for the category of the predicted “target” item. We used partial correlation to control for the mean evidence of the three non-target categories for each prediction (for similar approaches see Detre et al., 2013; Kim et al., 2014), and the pruning relationship between target evidence and subsequent memory held for the predicted items (Figure 5B). To rule out the possibility that memory for the mispredicted items was worse due to poor initial encoding of those items, we controlled for the perception strength of each item during encoding and found that the relationship between prediction strength and subsequent memory remained negative in the incongruent condition and absent in the congruent condition (Figure 5C). Finally, we ran an analysis to evaluate the specificity of the relationship between perception strength and subsequent memory. We used partial correlation to control for the amount of non-target category evidence during perception and confirmed that these relationships were specific to the evidence for the target category (Figure 5D).

These main analyses were also replicated with modified regressors to rule out possible confounds from varying ITIs and different times windows for prediction and perception. Fixed 3 s time windows from the onset of a trial were applied and then shifted 4 s forward to account for hemodynamic lag for both prediction and perception. Consistent with our original results, we found evidence of memory pruning only in the incongruent condition (β = −0.64, *P* = .004) and not in the congruent condition (β = −0.43, *P* = .087). The linear patterns of change in the perception-memory relationship also replicated, such that the relationship strengthened for the incongruent condition (β_lin_ = 0.51, *P* < .001) and weakened for the congruent condition (β_lin_ = −0.45, *P* = .089) as a function of repetitions (linear: β_inc_ = 0.51 vs. β_con_ = −0.45, *P* = .007).

### Mismatch signals in hippocampus

Linear regression analyses were applied to link mismatch signals in the hippocampus (Long et al., 2016) and prediction strength decoded from the ventral temporal cortex. The mismatch signal was defined as the average intensity of the signal during the *perception window*, and prediction strength was defined as the average classifier evidence during the *prediction window*. There was no reliable relationship between mismatch signals and prediction strength for either incongruent or congruent conditions (*Ps* > .05).

## DISCUSSION

When we reencounter a familiar situation, our brain anticipates what might come next based on previous encounters. This study demonstrates that the predictability of our experiences influences whether we prune memories of past events and also how we form memories of new ones. Our results show that when the temporal statistics across repeated experiences allow for the learning of abstract, conceptual relationships between events, this has two main consequences: (1) memory for past events is no longer impacted by prediction errors for new events, and (2) memory for new events declines because encoding becomes less focused on item-specific details.

We used fMRI pattern classifiers to track the explicit encoding and the covert prediction of visual stimuli during a continuous sequence of image presentations. The sequence was configured such that some of the stimuli (“cues”) repeated multiple times across the experiment. Cue stimuli were followed either by new items that belonged to different categories (incongruent condition) or new items that belonged to the same category (congruent condition). For cues in the incongruent condition, which afforded participants no reliable predictive information across repetitions, stronger neural evidence for a (incorrect) prediction – that the previous item to follow when the cue reappeared – were associated with worse subsequent memory for that mispredicted item. This replicates the memory pruning effect observed previously (Kim et al., 2014) that describes a form of error-driven statistical learning in which the memory trace for a mispredicted event is weakened, which leads to subsequent forgetting of that event. A key result of this study is that for cues in the congruent condition, which afforded participants the ability to learn across repetitions the category of item to expect, no memory pruning was observed. Furthermore, encoding of new items after the final appearance of congruent cues was impaired relative to incongruent cues: subsequent memory on average was worse for these congruent items, and the memory strength of individual items was unrelated to neural measures of their perception strength.

### Context-based predictions of items

Our neural analyses of automatic predictions relied on pattern classifier estimates for a “target” category following the repetition of a cue stimulus (e.g., classifier evidence for the target category of “animal” following the repetition of *cue* in the sequence “*cue,* badger*, …, cue, …*”). On average, the amount of classifier evidence following a cue repetition for the target category was not greater than for non-target categories (in this example: food, tools, vehicles). This was true for both incongruent cues and congruent cues, and by itself is inconsistent with the idea that cues triggered predictions of items from the previous repetition (see Results for the target selectivity from prediction betas). However, the results found from linking these neural measures of “prediction” on a trial-by-trial basis to subsequent memory outcomes suggests otherwise. For incongruent cues, stronger classifier evidence for the (predicted) target category was associated with weaker subsequent memory for those items (Figure 4A). This relationship held even after controlling for the amount of classifier evidence for the non-target categories at the time of prediction and for the initial perception strength of those target items from the previous repetition (Figure 5B and C). This robust link to item-specific behavioral outcomes is consistent with the interpretation that incongruent cues triggered specific predictions of previous items that were consequently pruned from memory after the prediction error.

Additional evidence for context-based predictions comes from the observation of *reduced* classifier decoding accuracy for the perception of the repeated cues compared to the single-exposure items (Figure 2B). Notably, this is the opposite relationship of classifier performance from within the localizer data alone, where the cue categories (face, scene) were decoded *better* than the other categories (animal, food, tool, and vehicle, Figure 2B). The elimination of the decoding advantage for the repeated cues could have arisen from two sources: first, from reduced processing of the now-familiar cues across repeated presentations (i.e., repetition suppression), and second, from the co-mingling of cue processing with an automatically triggered prediction for an item (or items) from another category, which would dilute the measurement of cue-specific neural activation. Repetition suppression has been linked to increased, not decreased, classifier evidence, which is believed to reflect a sharpening of the neural representation for repeated stimuli (Kim et al., 2014; Kok, Jehee, & de Lange, 2012). However, reduced variability in activity across voxels, which can occur due to repetition suppression, has also been associated with decreased classifier performance (Davis et al., 2014). Regardless of the cause, the reduction in cue decoding across repetitions cannot account for the key results linking prediction evidence for items and their subsequent memory.

The argument for item-specific predictions is further supported by the observation of a strong relationship between perception strength and memory for the final items in the incongruent condition. These items were remembered well, and their perception strength was directly related to their memory strength. This direct link between perception strength and memory suggests that individual item details were being encoded, which should in turn facilitate predictions of these items upon the next appearance of the cue. Note that it is likely that the encoding and subsequent prediction for these items would decrease if and when it was learned that these predictions were always violated, but this did not seem to occur in this study after only four repetitions.

### Context-based predictions of categories

On the other hand, in the congruent condition, cues were always followed by new items from a single category (e.g., a *cue* was always followed by an animal: “*cue*, badger, …, *cue*, tiger, …, *cue*, cow, …, *cue*, peacock”). It is possible that participants could learn this relationship for each cue and make explicit predictions at the category level. However, post-experiment questionnaires confirmed that, aside from participants noticing repeated presentation of the cues, they did not detect any structure in the order of stimulus presentations. Any differences in prediction between the experimental conditions would therefore be due to implicit learning of the transition probabilities associated with incongruent cues vs. congruent cues. It could be possible that stimulus-task associations were learned in the congruent condition (e.g., Anne Hathaway followed by a cooked/uncooked subcategorization task), rather than stimulus-stimulus association (e.g., Anne Hathaway followed by a food item). If the task itself were predicted from a given cue, it would have led to faster and more accurate performance for the congruent items. However, the encoding performance showed the opposite results in which the subcategory judgment for the congruent items was slower and less accurate than for the incongruent items (Figure 1A, main effect for item). These results suggest that there was no priming effect at the decision level for the task.

Similar to the results for incongruent cues, there was no direct neural evidence of prediction (target classifier evidence was not greater than non-target evidence) for congruent cues. Unlike incongruent cues, however, there was no relationship between the neural evidence for the “predicted” item (i.e., the item that followed the cue on its prior appearance) and that item’s subsequent memory (Figure 4B). These data do not support an inference that participants were making item-specific predictions for congruent cues. Rather, they suggest that congruent cues either (1) triggered no predictions at all, or (2) triggered predictions at the *category* level rather than at the *item* level (e.g., “expect some animal” instead of “expect that badger”). Two additional results provide converging evidence for the latter interpretation. First, if predictions were categorical, this should focus attention on categorical information, rather than item-specific details, during the encoding of new items, which in turn should lead to worse subsequent memory for these items. Indeed, we found that stronger prediction evidence after a congruent cue was associated with worse subsequent memory for the next item, which by design was a new exemplar of that category (β = −0.64, *P* = .008). This relationship between prediction strength and memory for new items was not true for incongruent cues (β = 0.01, *P* = .47). Second, as reported in Figure 4B, the perception strength for new items following congruent cues became decoupled from their subsequent memory after the second appearance of the cues.

Together, we believe these results are most consistent with the idea that congruent cues triggered categorical predictions rather than no predictions at all. The congruent condition results reflect the consequences of statistical learning focused not on item-specific details, but rather on abstract conceptual information (Brady & Oliva, 2008). Across multiple experiences, overlapping features (i.e., the semantic category) of individual events were extracted to form more generalized knowledge about these specific situations, similar to the formation of memory schemas (O’Reilly & Norman, 2002; Tse et al., 2007; Marlieke T. R. van Kesteren, Brown, & Wagner, 2016; Marlieke T R van Kesteren et al., 2012). Once this general inference was developed, this focused future encoding on category-level information and the specific episodic details of new items were forgotten (Marlieke T. R. van Kesteren et al., 2016; Marlieke T R van Kesteren et al., 2012).

Our analyses relied on category-specific fMRI pattern classifiers to covertly measure implicit predictions of previous items from repeated contexts. Category classifiers can produce more robust decoding than sub-category classifiers or item-level classifiers, but of course they lack item specificity. We decided against proceeding with an item-level decoding approach due to insufficient classifier sensitivity in early pilot data. Instead we relied on category information and therefore could not distinguish predictions for individual exemplars of a category from generic category predictions.

Instead, we leveraged the relationship (or lack thereof) between these category-specific neural estimates and the item-specific behavioral measures for each stimulus to speculate on the nature of these predictions. Future work should use neural analyses sensitive to item-level representations (e.g., representational similarity analysis; Kriegeskorte, Mur, & Bandettini, 2008) to more directly test this idea.

### Memory pruning

Results for the incongruent condition in the present study, with cues followed by novel items from novel categories, were consistent with previous findings on memory pruning (Kim et al., 2014) which found a negative relationship between neural prediction strength for mispredicted items in a temporal sequence and subsequent memory for those items. Memory pruning is a form of error-driven learning that is consistent with predictions of the non-monotonic plasticity hypothesis (NMPH) (Detre et al., 2013; Lewis-Peacock & Norman, 2014a; Newman & Norman, 2010) which claims that moderately activated memories can lead to weakening and subsequent forgetting of those memories. Here, moderately active memories were created by the automatic context-based predictions that occurred following cue repetitions in the incidental encoding task. In Figure 3A, the *prediction strength* for the predicted items following a repeated cue is contrasted with the *perception strength* for the items that actually appeared. As would be expected, the distributions of classifier evidence values show that perception strength (*M* = 0.70) was reliably higher than prediction strength in both conditions (*M* = 0.52, *Ps* < .001). Taking classifier evidence as an index of the strength of “memory activation”, we see that prediction leads to more moderately active representations (compared to perception), and the NMPH predicts that these memory representations would be more vulnerable to weakening and long-term forgetting. In a separate whole-brain searchlight analysis (Figure 6), we found that the prediction-based pruning effect for the incongruent condition was largely clustered in the VTC (10.1 % of all identified voxels). The identified voxels in the VTC were 31.4% of all voxels in this region, which is a substantial concentration given that we used M = 26.6% voxels for classification through feature selection in the main analysis. It is apparent from this analysis that the memory pruning results found in VTC were also found in a distributed network of voxels in parietal and frontal cortex. Whether these regions operate in parallel or in some other coordinated fashion to support memory pruning is an interesting topic for further research.

According to this framework, predictions of a specific event are realized by anticipatory activation of its memory representation, resulting in relatively weak activation of its memory trace (compared to activation during the initial perception of the event). If this prediction is confirmed, its neural activation will increase, as will the strength of its representation in long-term memory, due to additional processing of the event. If the prediction is violated, its reactivated memory representation will not be enhanced beyond this moderately activated state which, according to the hypothesis, can lead to weakening and forgetting of the item. In temporal contexts that allow for more abstract statistical learning (e.g., our congruent condition), the predictions may not contain representations of specific events from the past (e.g., “some animal” might be expected but not “that badger”). The confirmation of categorical predictions in the congruent condition should not then invoke NMPH-based memory weakening processes for those predictions. The lack of specificity in these predictions may have the effect of shielding the memories of those specific events from modification.

### Memory pruning vs. memory integration

It could be argued that cues in the incongruent condition should trigger memory integration (Greve, Abdulrahman, & Henson, 2018; Morton et al., 2017; Preston & Eichenbaum, 2013; Schlichting & Frankland, 2017; Schlichting et al., 2015; Schlichting & Preston, 2015; Zeithamova, Dominick, et al., 2012; Zeithamova, Schlichting, et al., 2012) rather than memory pruning, such that all items maintain their associations to the repeated context, similar to inferential learning (Morton et al., 2017; Preston & Eichenbaum, 2013; Schlichting & Frankland, 2017; Schlichting et al., 2015; Schlichting & Preston, 2015; Zeithamova, Dominick, et al., 2012; Zeithamova, Schlichting, et al., 2012). Recent evidence suggested that prediction error weakens overlapping representation between the mispredicted item and its context on the predicted item, leading to differentiation of their neural patterns in the hippocampus (Kim et al., 2017). However, the same neural consequences have also be observed for memory integration (Zeithamova, Dominick, et al., 2012). The hippocampus has a critical role, not only for memory integration (Eichenbaum, 2000; Marlieke T. R. van Kesteren et al., 2016) but also in mismatch detection (Kumaran & Maguire, 2007; Long et al., 2016) for prediction errors. In a recent study by Long and colleagues (Long et al., 2016), the activation of the hippocampus was found to be positively correlated with prediction errors, and even more so if the mispredicted item was semantically related to the actual item. (This is similar to the congruent condition in the present study.) The hippocampus was not recruited when predictions were correct or unrelated semantically to the novel events. This suggests that mismatch signals are key for triggering updating of existing memories (Kumaran & Maguire, 2007; Long et al., 2016; Schlichting & Preston, 2015). In our data, however, we were unable to find any relationship between prediction strength and hippocampal activation, or any evidence of hippocampal involvement when mismatched predictions were semantically related to the new events (i.e., in the congruent condition). Unlike Long and colleagues’ study or inferential learning studies (Schlichting et al., 2015; Zeithamova, Schlichting, et al., 2012) in which the participants explicitly learned word-picture or picture-picture associations in the pre-training phase, the cue-item associations were implicitly learned in our study. Explicit predictions based on over-learned associations might be too strong to trigger pruning of existing memories, but rather may promote integration of the new semantically related items (Morton et al., 2017; Schlichting & Preston, 2015).

We could not find hippocampus involvement, such that the prediction signals would be reflected through “pattern completion” after cue re-appearance (Kim et al., 2017). This suggests that the hippocampal signals would be detectible only detectible when association learning is more stable, e.g., via explicit association or multiple exposures of implicit association.

### Less robust memory for items that occur in predictable contexts

In our study, the congruent condition involved repeated visual cues that were consistently followed by items from a single category. Results suggest that participants implicitly learned these relationships, as both behavioral evidence and neural evidence in this condition diverged from the incongruent condition in which the cues were always followed by a new item from a new category. Specifically, memory for the final item in the congruent contexts was worse than previous items in those same contexts (Figure 1B), and there was no relationship between neural evidence of perception for these items and their subsequent memory strength (Figure 4B). Together these results suggest that in temporal contexts with greater predictability (e.g., when the category of the next item can be anticipated), encoding of new items may be reduced (but see Friedman, 1979; Gronau & Shachar, 2015; van Kesteren et al., 2010; Zwaan & Radvansky, 1998). Consistent with this idea, Rommers and Federmeier (2018) recently demonstrated evidence that predictable information is processed more weakly than unpredictable information. When words reappeared in predictable contexts, the neural responses measured using electroencephalography (EEG) indicated that the words were processed less than repeated words in unpredictable contexts. Specifically, the repetition priming effects were diminished in the N400 and LPC components of the EEG signal. From these results, the authors suggested that predictability allows the brain to operate in a top-down “verification mode” at the expense of detailed stimulus processing. Our data are consistent with these findings in language processing, and they extend the idea that predictability of perceptual events (here, in sequences of visual objects) leads to reduced encoding of new stimuli.

## Conclusions

The learning processes observed in this study are examples of adaptive forgetting that allow for the efficient use of the brain’s memory systems (Kim et al., 2014, 2017; Lewis-Peacock & Norman, 2014a; Wylie, Foxe, & Taylor, 2008). Being able to anticipate the demands required of us in familiar situations can help us to respond more effectively and proactively. Pruning unreliable memories via statistical learning mechanisms supports this behavior by reducing interference during context-based retrieval of relevant memories (Kumaran & Maguire, 2007). Here, we demonstrated that the stability of associative memories is evaluated over multiple exposures to the context in which those memories were acquired. When a context afforded no accurate predictions, previous experiences were nonetheless anticipated, perhaps reflecting a persistent, but futile, effort to learn the statistics of the environment. Memory for these experiences was pruned when their predictions were violated. When a context afforded general information (but not specific details) about what to expect, the previously encountered events were no longer predicted, and their memories were shielded from pruning. However, new learning was also diminished in these more predictable contexts. These findings deepen our understanding of how associative memories are formed and updated by demonstrating that our ability to predict the future influences how we remember the past.

